# Accelerating Insight Discovery in Large Biomedical Text with Scalable Processing Framework

**DOI:** 10.1101/2025.08.14.670384

**Authors:** Dongeun Kim, Megan Hauptman, Matthew T. Patrick

## Abstract

Large language models are increasingly being used by dermatology professionals to support diagnostic investigation, patient education, and medical research. While these models can help manage information overload and improve efficiency, concerns persist regarding their accuracy and potential reliance on dubious sources. We introduce Quanta, a hybrid system that combines large language models with established evaluation metrics, such as cosine similarity, to enable efficient summarization and interpretation of curated research corpora. This methodology ensures that synthesized insights remain domain-specific and contextually relevant, thereby supporting clinicians and researchers in navigating the expanding digital landscape of dermatology literature. Deployed within an interactive chatbot, the tool delivers direct answers to user queries, provides cross-publication insights, and can suggest new directions for research. Comparative evaluations on benchmark datasets demonstrate improvements in accuracy, efficiency, and computational cost, with the curated document approach enhancing reliability and reducing misinformation risk.

## Introduction

Large language models (LLMs) are a form of deep neural network, which harness advanced computational techniques and massive training data, to provide meaningful output in response to textual queries (Kasneci et al., 2023). LLM usage has increased dramatically in recent years. A survey from the American Academy of Dermatology found more than half of dermatologists use LLMs every week (Gui et al., 2024), with the most popular application being to investigate diagnoses and medical management. Further uses include preparing patient handouts and assisting in dermatology research (Jin and Dobry, 2023). LLMs have the potential to enable dermatology researchers and clinicians to keep up to date with a rapidly increasing body of knowledge. They address information overload and facilitate greater efficiencies in work, by handling laborious and time-consuming tasks, particularly those that involve the extraction of meaningful insights from massive text corpora. However, it is important to consider these benefits in the context of any introduced risks.

The vast range of data on which LLMs are trained can make it possible for irrelevant or disreputable sources to affect the results (De Cassai and Dost, 2024, Lechien et al., 2024). Furthermore, the influence of these sources can be difficult to perceive, because of the complicated and unclear ways in which LLMs select and summarize information, often combining multiple documents together. Many dermatologists are concerned about the accuracy of LLMs (Gui et al., 2024), particularly since mistakes could have a negative impact on patients’ health. A study into using an LLM for novel idea generation in research also found the feasibility of these ideas (evaluated by expert review) was lower than those identified by humans (Si et al., 2024). Recognizing that commonly used LLMs may include unreliable or irrelevant sources, users benefit from providing a specific set of curated and trusted literature on which to ask their questions (Ben Abacha and Demner-Fushman, 2019). However, even powerful language models, such as GPT-4 and other currently available tools, are limited in the number and/or size of files that can be analyzed (Wang et al., 2024). The need for effective methods to process and analyze vast amounts of textual information is becoming increasingly important, as digital resources grow in number and accessibility. We aim to address these problems by introducing a new tool (Quanta), which can efficiently process and synthesize relevant information from large-scale textual data.

To extract meaningful insights from heterogeneous textual data, researchers have explored various methods, including traditional lexical similarity measures such as Cosine Similarity, Jaccard Similarity, and Term Frequency - Inverse Document Frequency (TF-IDF) (Rinjeni et al., 2024). Cosine similarity is particularly effective for tasks such as finding related information (semantic search), clustering similar items, and retrieving relevant documents. It takes advantage of multiple distinct semantic dimensions to capture different aspects of meaning and, as a result, cosine similarity is well-suited for analyzing text in a way that preserves and reveals the underlying meaning of the output.

However, traditional similarity approaches struggle to fully capture semantic meaning, particularly in more contextually rich documents, as they lack mechanisms for modeling word order and context. So, more advanced deep learning models have been proposed, including Bidirectional Encoder Representations from Transformers (BERT) (Gorenstein et al., 2024). These approaches take advantage of multiple layers of abstraction to learn numerical (vector) representations of each text, embedding their semantic meaning in a way that can be effectively used for computation. Yet, deep learning-based methods can be computationally expensive, particularly when handling vast datasets. To address these limitations, this research introduces a hybrid methodology that combines conventional similarity measures (e.g. cosine similarity) with transformer-based semantic models such as BERT.

A key innovation of our approach is the integration of LangChain^1^ to optimize query decomposition, dynamic prompt generation, and contextual refinement. By abstracting much of the implementation complexity, LangChain enables seamless integration of recent language technologies without requiring extensive low-level programming. Specifically, it supports the transformation of raw text into numerical representations suitable for semantic search, manages retrieval operations from the database of vector embeddings, and ensures that the input provided to the language model follows a consistent and structured format. This foundation contributes to a modular and scalable system design, allowing for consistently accurate and context-aware responses across diverse clinical and biomedical text sources.

By allowing dermatologists and dermatology researchers to input their own carefully curated and pertinent set of literature into the system, we ensure that the synthesized information remains domain-specific and contextually relevant, while delegating the time-consuming tasks of document review to the LLM. Furthermore, we leverage ChromaDB for efficient embedding storage and fast nearest-neighbor retrieval, especially considering the usage of LangChain, which natively supports ChromaDB for efficient storage of the vector database. This integration simplifies retrieval workflows by enabling seamless coordination between embedding generation, metadata filtering, and document ranking, without the need for manual configuration or complex indexing.

## Results

### Overview of Approach

We propose an integrated pipeline built upon recent technologies, as illustrated in **(Figure 1)**.

**Figure 1.**
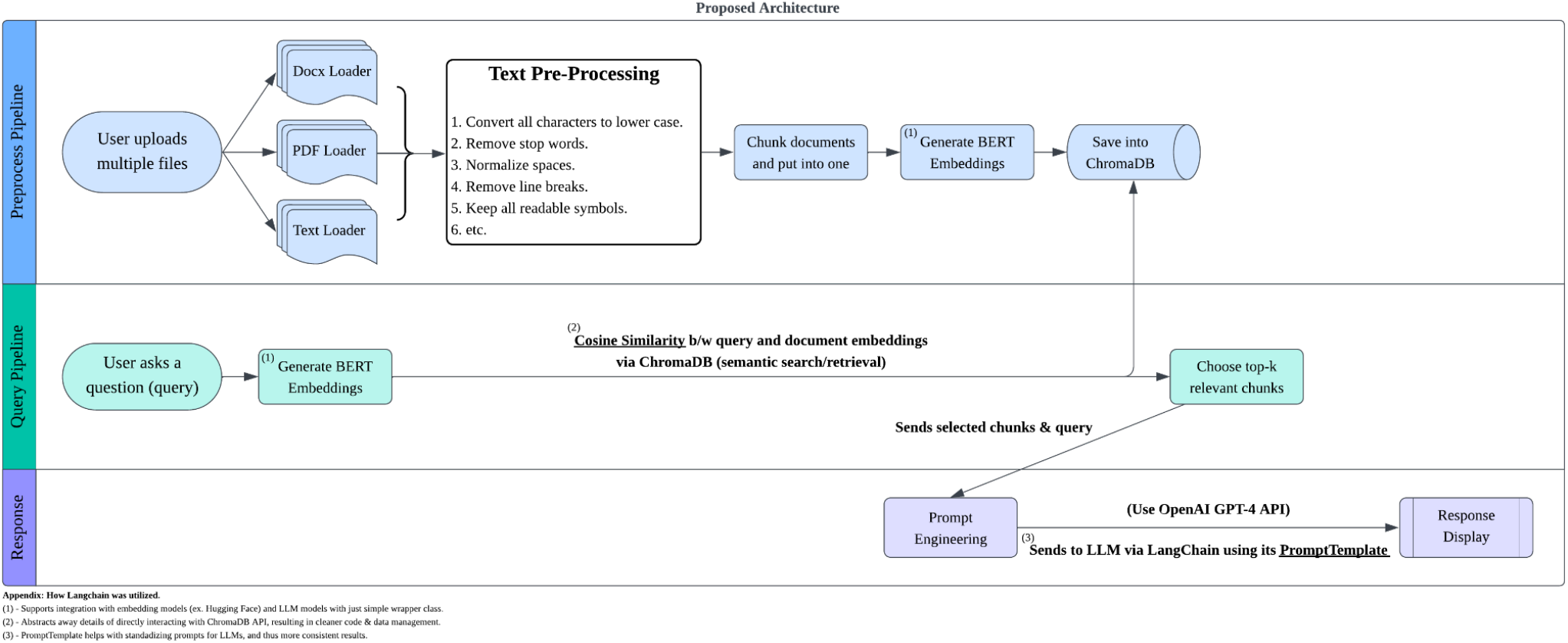
Quanta Architecture. Three-stage architecture designed to enable intelligent querying over user-uploaded documents. Consists of the Preprocess Pipeline, Query Pipeline, and Response Pipeline, integrating BERT embeddings, ChromaDB, and OpenAI’s GPT-4 through LangChain.

The proposed architecture is designed to enable semantic search and response generation over heterogeneous document formats using a modular, retrieval-augmented generation (RAG) framework. The system is composed of three main pipelines, which interact via shared vector representations and LangChain integration: Preprocess Pipeline, Query Pipeline, and Response Pipeline.

### Preprocess Pipeline

The preprocess pipeline is responsible for preparing the uploaded documents for semantic indexing. Users can upload documents in various formats (DOCX, PDF, TXT), which are parsed using appropriate loaders (DocxLoader, PDFLoader, TextLoader, etc.). Then, the text is preprocessed to ensure consistency and reduce noise, by applying a series of normalization operations. These include conversion to lowercase, removal of stop words and line breaks, as well as normalization of spacing. The processed texts are then divided into semantically manageable chunks. Each chunk is embedded using a BERT-based encoder to produce fixed-size vector representations. The resulting embeddings are stored in ChromaDB, a vector database optimized for fast similarity search when used with LangChain.

### Query Pipeline

The query pipeline processes incoming user questions and retrieves the most semantically relevant document chunks by applying content relevance filtering. This is achieved by embedding user queries using the BERT model, to ensure consistency in semantic space with the BERT-embedded document chunks. These embeddings, from user queries and documents, are then used to compute cosine similarity, and the top *k (5 by default, but can be adjusted)* most similar chunks are selected as context.

### Response Pipeline

The response pipeline formulates a natural language response based on retrieved document content. A system prompt (**Supplementary Note 1**) is formatted using LangChain’s PromptTemplate functionality and provided to OpenAI’s GPT-4 language model^2^, along with the selected chunks and original query, to ensure the response is well-structured, coherent and context-aware. This is possible as it only focuses on the most relevant contents, which is filtered and selected during the query pipeline. The output of each query is displayed via the Quanta interface, which was created using Streamlit^3^ **(Figure 2)**. LangChain plays a central role in helping us to simplify implementation complexities, as follows: (1) Embedding Integration: LangChain supports wrapper classes for generating embeddings (e.g., via Hugging Face), (2) Retrieval Abstraction: LangChain simplifies ChromaDB access, eliminating the need for direct application programming interface (API) interactions, (3) Prompt Standardization: The PromptTemplate module ensures consistency and quality in the prompts passed to the LLM.

**Figure 2.**
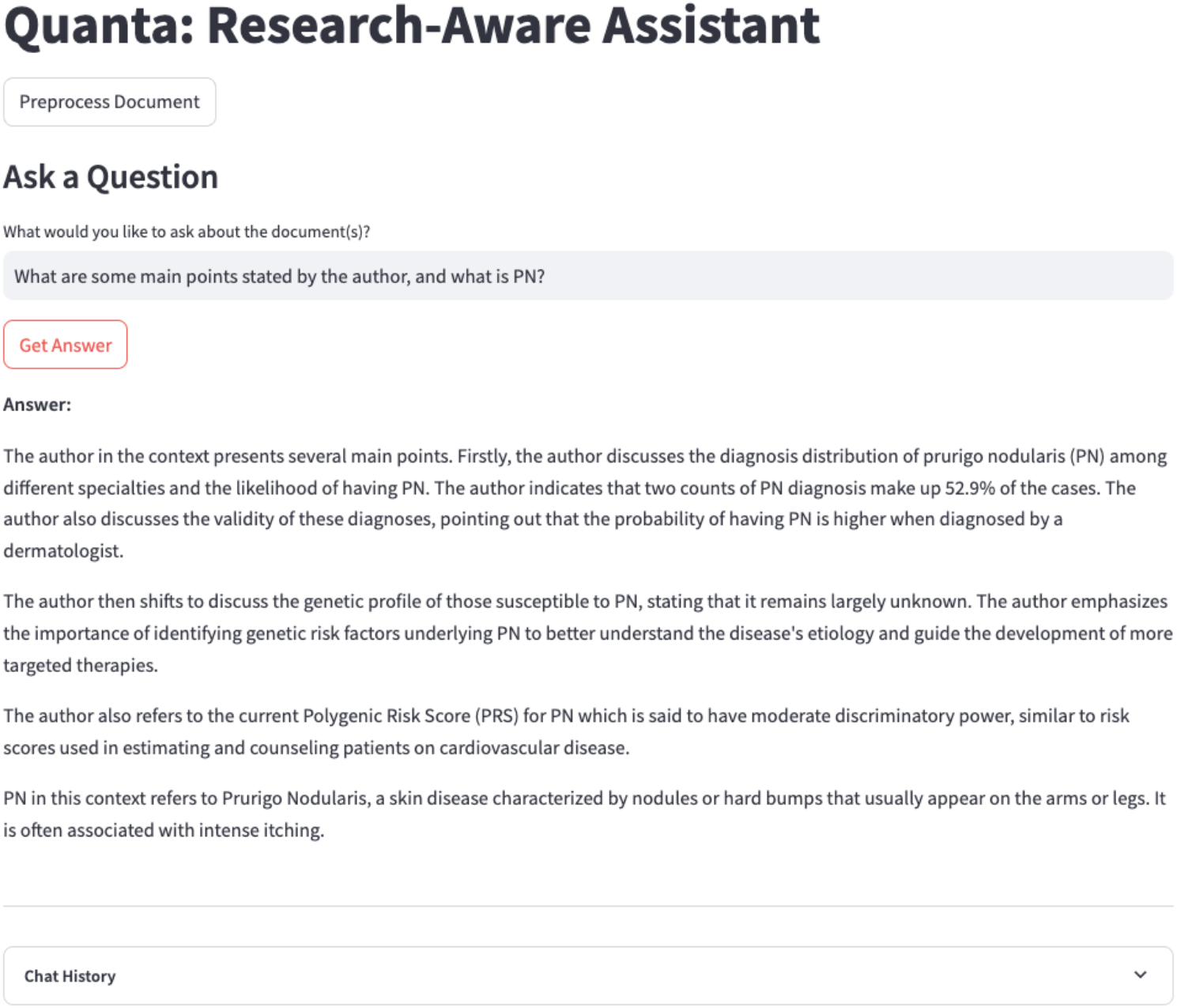
Interface for Quanta Assistant.

### Embedding Model Selection and Sensitivity Analysis

We evaluated different configurations for semantic search and information retrieval, using the Benchmarking Information Retrieval (BEIR) framework^4^. It comprises 19 datasets for retrieval tasks across a wide range of domains, including biomedical, finance, and open-domain question answering. They each have comprehensive “ground-truth” labeling that facilitates the reliable comparison of embedding methodologies.

Using 5 different embedding models (see **Materials and Methods**), **Figure 3** presents our comparison of the Euclidean L2 (blue) and cosine similarity (orange) approach for semantic search and information retrieval performance. The cosine similarity approach consistently outperformed L2 across all metrics (**Table 3**). When differences were present, the cosine approach achieved approximately 48.7% to 76.9% higher mean scores for Normalized Discounted Cumulative Gain (NDCG), around 7.1% higher for F1-score, roughly 33.3% higher for Recall@5, and between 20% and 100% higher for Mean Reciprocal Rank (MRR). It also demonstrated shorter average processing time, decreasing the duration by 95.6% compared to the L2 approach on average across various embedding models. This performance advantage stems from cosine similarity’s ability to capture semantic relevance more effectively by emphasizing the direction of embeddings rather than their magnitude, making it better suited for high-dimensional spaces where vector orientation encodes core semantic meaning.

**Figure 3.**
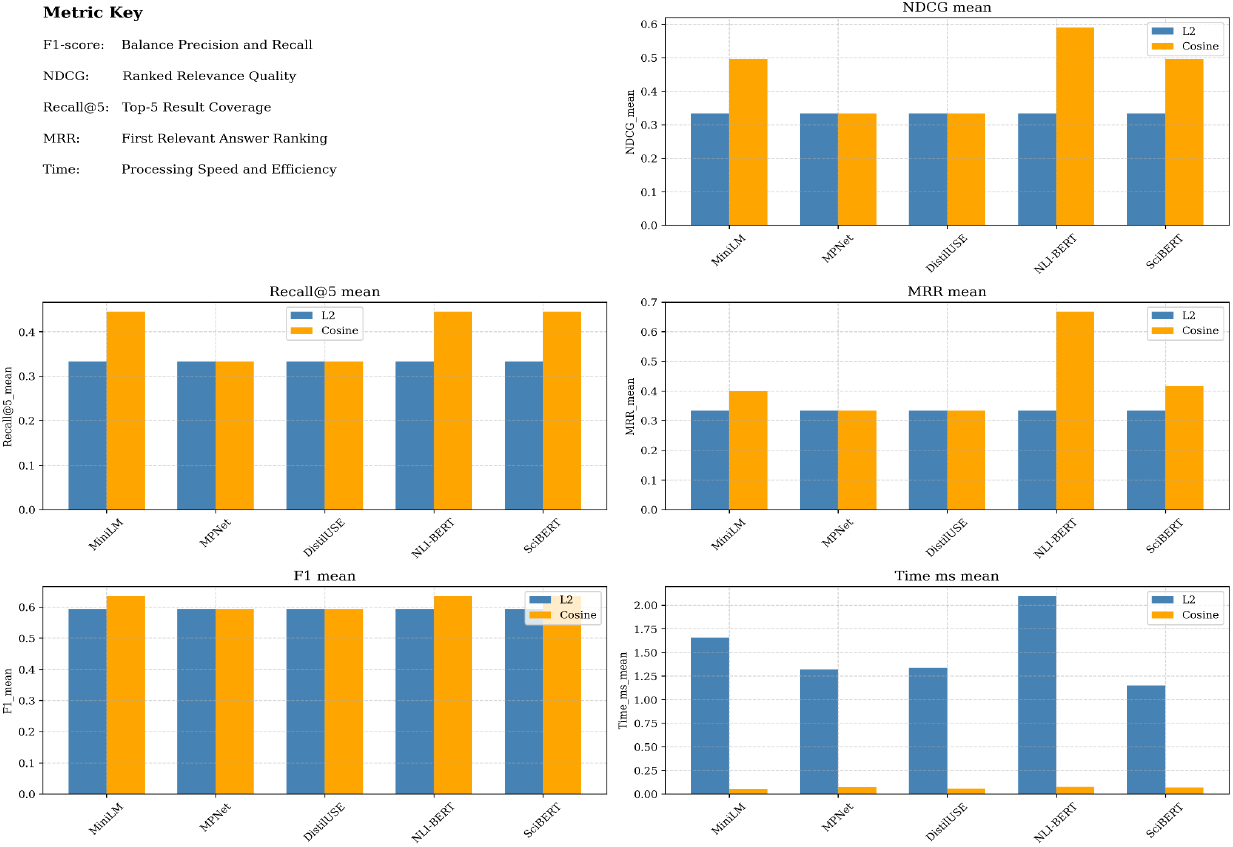
Comparative performance of cosine vs. L2 similarity across embedding models. Retrieval performance of various embedding models (MiniLM, MPNet, DistilUSE, NLI-BERT, SciBERT) when evaluated using five key metrics: F1-score, NDCG, MRR, Recall@5, and computational time.

Of the embedding models we evaluated, NLI-BERT performed as well as or better than all other models across all metrics (**Figure 3**), with highest mean scores for NDCG, MRR, and shorter average processing time, suggesting it to be an optimal embedding model for the Quanta retrieval pipeline. **Figure 4** presents a heatmap of standard deviations for the retrieval metrics scores of each embedding model. MiniLM had the lowest average standard deviation at 0.342. However, this is just 4.8% lower than the 3rd best model, NLI-BERT (0.358), indicating only a slight difference in retrieval consistency. Given this minimal variance gap, the combination of consistent high mean scores **(Figure 3**) and low variance makes NLI-BERT a strong candidate for robust semantic search at large-scale.

**Figure 4.**
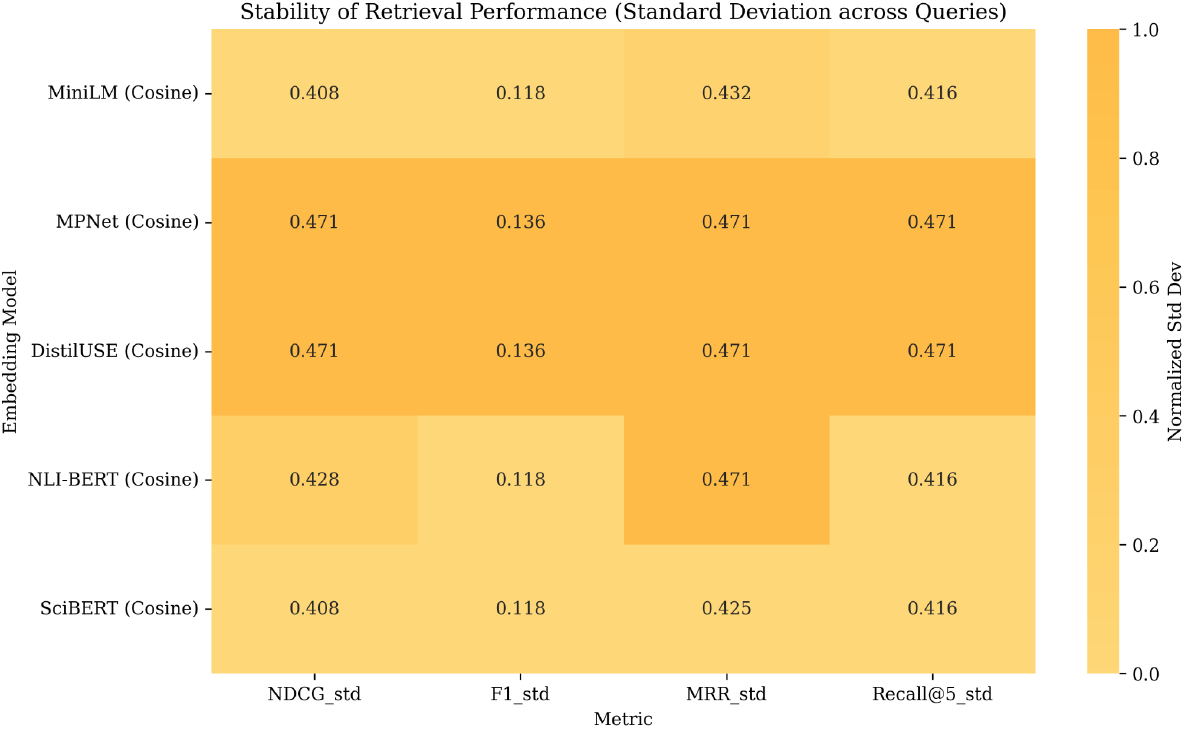
Heatmap of standard deviations across metrics for each embedding model. Visualizes the stability of each tested embedding model by showing the standard deviation across retrieval metrics scores. Lower standard deviation values indicate greater consistency, shown by lighter opacity of yellow.

### Validating Integration of LangChain

We then tested whether integrating LangChain into our vector-based retrieval pipeline could enhance semantic matching performance. This was evaluated in terms of the relevance of retrieved answers using BEIR, compared to raw vector similarity alone (**Tables 1** and **2)**. For each metric, we conducted paired t-tests to assess the distribution of differences across query-level scores. The paired t-test results showed statistically significant improvements for LangChain, validating our hypothesis with p-values of 0.01302 (F1), 0.00920 (NDCG) and 0.01116 (Recall@5), respectively. Similarly, a Wilcoxon signed-rank test for MRR confirmed a significant effect, with a p-value of 0.01776, further supporting the hypothesis that LangChain improves semantic retrieval performance.

**Table 1.** Baseline BERT Results. *SciBERT was used as our Baseline BERT, as it showed best performance in Figure 3 and Figure 4.

| query_id | NDCG | F1 | MRR | Recall@5 | Time(ms) | Memory Bytes |
| --- | --- | --- | --- | --- | --- | --- |
| PLAIN-2 | 0.0 | 0.493 | 0.0 | 0.0 | 11.956 | 3754374 |
| PLAIN-23 | 0.0 | 0.493 | 0.0 | 0.0 | 2.037 | 3730572 |
| PLAIN-78 | 0.399 | 0.605 | 0.1 | 0.0 | 1.996 | 3730572 |
| PLAIN-307 | 0.813 | 0.662 | 1.0 | 0.667 | 2.796 | 3730572 |
| PLAIN-332 | 0.347 | 0.552 | 0.111 | 0.0 | 2.2 | 3730572 |
| PLAIN-371 | 0.0 | 0.493 | 0.0 | 0.0 | 2.496 | 3730572 |
| PLAIN-430 | 0.399 | 0.599 | 0.143 | 0.0 | 2.031 | 3730572 |
| PLAIN-468 | 0.0 | 0.493 | 0.0 | 0.0 | 1.931 | 3730572 |
| PLAIN-499 | 0.0 | 0.488 | 0.0 | 0.0 | 2.018 | 3730572 |
| PLAIN-531 | 0.399 | 0.542 | 0.167 | 0.0 | 1.933 | 3730572 |

*Used the best performing embedding models based on Graph 1.*

Furthermore, a graphical representation of the per-query performance scores is shown in the violin plot in **Figure 5**, which emphasizes both the distribution shape and median values for each metric. In general, the LangChain system (orange) exhibits a higher median across most metrics compared to the baseline system (blue), indicating better performance. Notably, the NDCG (ranked relevance quality) distribution is skewed substantially higher for LangChain, reflecting stronger retrieval precision, where relevant items were frequently placed near the top of the ranked output. **Figure 5**’s visualization complements the statistical findings by offering a more intuitive view of LangChain’s relative effectiveness. Finally, numerical values from **Table 1 and Table 2** show that although LangChain-based retrieval incurred higher latency due to deeper semantic processing, it reduced memory usage by approximately 96.98% (see **Supplementary Note 2** for calculation details), significantly improving efficiency in storage and scalability.

**Table 2.** LangChain Results.

| query_id | NDCG | F1 | MRR | Recall@5 | Time(ms) | Memory Bytes |
| --- | --- | --- | --- | --- | --- | --- |
| PLAIN-2 | 1.0 | 0.784 | 1.0 | 1.0 | 141.661 | 525838 |
| PLAIN-23 | 0.0 | 0.497 | 0.0 | 0.0 | 84.559 | 128249 |
| PLAIN-78 | 0.487 | 0.640 | 0.2 | 0.333 | 87.023 | 105885 |
| PLAIN-307 | 1.0 | 0.874 | 1.0 | 1.0 | 25.891 | 107519 |
| PLAIN-332 | 1.0 | 0.640 | 1.0 | 0.5 | 69.783 | 106922 |
| PLAIN-371 | 0.0 | 0.497 | 0.0 | 0.0 | 71.108 | 112043 |
| PLAIN-430 | 1.0 | 0.720 | 1.0 | 0.5 | 22.643 | 104288 |
| PLAIN-468 | 0.487 | 0.665 | 0.333 | 1.0 | 67.364 | 103120 |
| PLAIN-499 | 1.0 | 0.547 | 1.0 | 0.077 | 77.661 | 101681 |
| PLAIN-531 | 0.487 | 0.535 | 0.333 | 0.059 | 73.760 | 105599 |

*Additional noise intentionally added to simulate real-world variation and test LangChain's robustness.*

**Figure 5.**
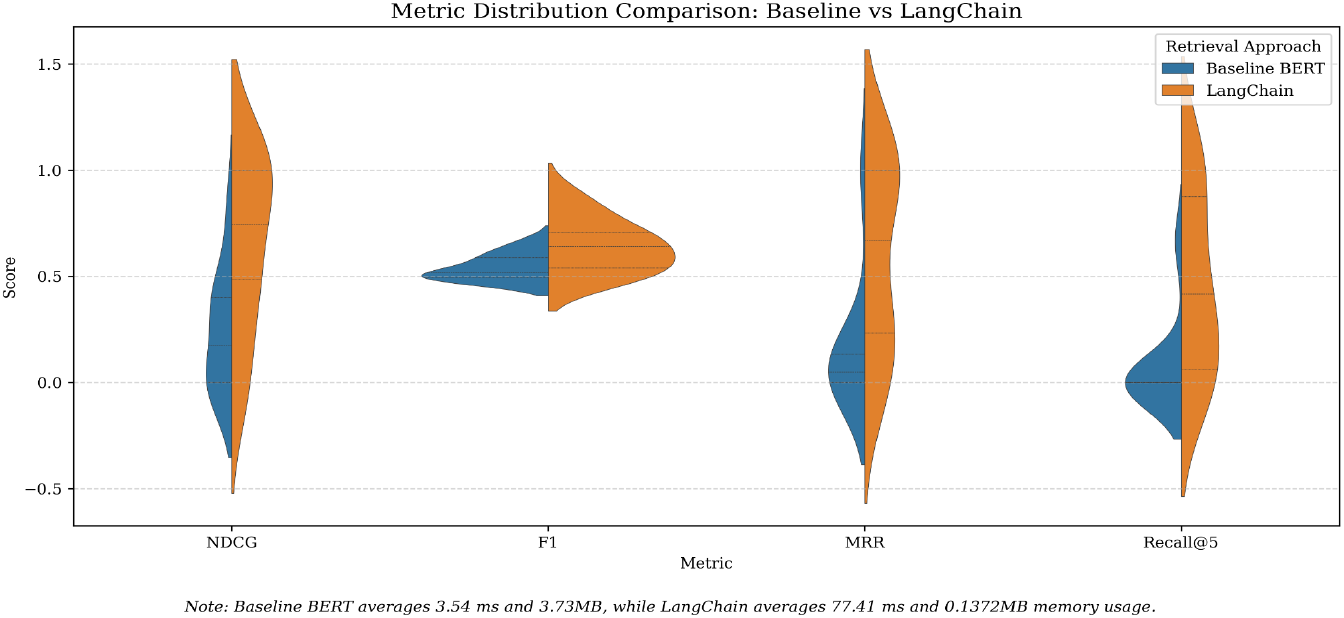
Violin plot of LangChain-integrated and baseline retrieval performances. Visualizes distribution of per-query F1-scores between the baseline vector retrieval pipeline and the LangChain-enhanced retrieval system. Each distribution reflects the spread and median value of the results across 50 test queries.

### Assessing Output

To validate Quanta’s abilities in a real-world setting, we tested whether it could accurately locate answers from a curated set of documents. The focus was to determine if our model could retrieve correct responses when asked specific questions. After uploading six key papers on prurigo nodularis, chronic itch, and cutaneous gene dysregulation, we asked a series of research-relevant questions. The responses (**Supplementary Note 3**) highlight the ability of our tool to provide meaningful insights, rooted in the literature, that may not be immediately obvious to the user. This demonstrates how our approach could be used to help facilitate further research, or provide valuable information in a clinical setting.

## Discussion

By combining advanced modeling techniques (large language models, LangChain and ChromaDB) with carefully tested evaluation techniques, including cosine similarity, we have developed an effective new tool for summarizing and interpreting large, curated sets of literature. Our approach can be of great use to clinicians and researchers by reducing the cognitive and time burden of manually reviewing extensive research papers, enhancing access to domain-specific knowledge, and facilitating discovery of nuanced insights that may span across multiple publications. As our model is incorporated into an interactive chatbot system that dynamically retrieves information from research papers, it can provide users direct responses to specific questions, as well as generate new research topics.

Our system precomputes document embeddings using OpenAI models and stores them in an internal document structure in memory, where cosine similarity is used to rank relevance at query time. This design allows fast retrieval, making it well-suited for interactive use cases. Furthermore, our hybrid retrieval strategy (combining dense vector similarity with metadata-based filtering) narrows down the search space before full ranking is computed, thus significantly improving our efficiency. Finally, our architecture supports content truncation and refinement, allowing us to flexibly limit the input length sent to the LLM while maintaining semantic completeness. We conducted a comparative analysis of our modeling decisions on the well-established BEIR datasets, evaluating performance in terms of accuracy, efficiency, and computational cost. Our results indicate that the model we developed can improve the synthesis of insights from large-scale research datasets.

Nevertheless, alternative approaches have been developed, and it is important to consider our work in the context of previous research. Hooper et al. (Hooper et al., 2024) introduced a method (Squeezed Attention) to accelerate long-context LLM inference by clustering fixed prompt keys based on semantic similarity and selectively computing attention only over the most relevant clusters. This achieves substantial improvements in speed and memory efficiency particularly for applications with long prompts; however, it requires inputs to be constrained to fixed contexts. By contrast, our approach addresses the challenge of big data by embedding content into a vector space and combining it with metadata-based filtering. This allows our approach to be more flexible, providing real-time semantic retrieval across dynamic and heterogeneous user queries for a diverse range of documents and tasks.

Other implementation decisions we could have made, to address the challenges of extracting meaningful insights from massive text corpora, include the combination of cosine similarity and principal component analysis (PCA). To improve computational efficiency and reduce memory usage, cosine similarity is often used in conjunction with PCA, which compresses high-dimensional embedding vectors by projecting them onto a lower-dimensional subspace. PCA does this by identifying the directions (principal components) along which the data varies the most and retaining only the top components that capture the greatest variance. This effectively removes less informative or redundant dimensions. Consequently, efficiency is gained at the cost of semantic nuance, which can negatively impact both the accuracy and relevance of retrieved responses.

Another approach is the Map-Reduce Summarization Method^5^, a pipeline designed to overcome context-length limitations. This method involves segmenting long documents into manageable chunks, summarizing each chunk individually while calculating token constraints, and then synthesizing these intermediate summaries into a final condensed summary. By summarizing in stages, this method enables the system to handle lengthy documents that would otherwise exceed input limits for large language models. While this approach effectively mitigates context window constraints, it introduces notable limitations. First, it leans toward summary-based retrieval rather than enabling access to the original full-text content, potentially omitting important contextual information. Second, because the summaries are generated independently of specific queries, the resulting outputs tend to be generalized and may lack the specificity required for rigorous research inquiries. Finally, this approach often incurs significant computational and financial costs due to repeated external API calls throughout the summarization pipeline.

Nevertheless, there are some limitations of our work that future research could investigate. LLMs are known to exhibit hallucinations (Huang et al., 2025), generating confident but inaccurate responses that can compromise the reliability of downstream research or clinical applications. Prompt engineering strategies, including techniques such as Chain-of-Thought (CoT) prompting (Wei et al., 2022), which encourages the model to reason step-by-step before producing an answer, might offer a way to reduce hallucinations. In addition, other plausible approach could be citation-constrained prompting, where the model is explicitly required to reference specific uploaded documents as the basis for its responses, else return “insufficient evidence” if no source is found.

A key advantage of our framework lies in its use of a curated and controlled set of documents, a setup that not only reduces the likelihood of misinformation but also gives researchers confidence in the trustworthiness of the source material. Moving forward, we aim to expand this approach by incorporating optional domain-specific filters (ex. restricting retrieval to clinical studies or expert-reviewed content for biomedical setting). This will further strengthen the system’s value as a transparent tool for navigating large-scale scientific literature.

## Materials & Methods

To enhance performance and ensure an optimal embedding model tailored for large-scale dermatological research applications, a structured evaluation process was implemented. We assessed the effectiveness of various embedding models through retrieval accuracy and stability, thereby identifying optimal implementation decisions for the Quanta model, as well as confirming the advantage of using cosine similarity over the Euclidean (L2) method, and investigating the benefits of LangChain integration.

### Computational Setup

The testing procedure was performed on standard computational hardware without the need for specialized external equipment. Yet, to ensure optimal performance and minimal latency, it is recommended to utilize a computer with at least 8GB of RAM.

### Selection of Embedding Models

Five widely adopted embedding models were selected based on their popularity, computational efficiency, and effectiveness in capturing semantic meaning: MiniLM (Wang et al., 2020), Masked and Permuted Pre-training for Language Understanding (MPNet) (Song et al., 2020), DistilUniversal Sentence Encoder (DistilUSE) (Reimers and Gurevych, 2019), Natural Language Inference BERT (NLI-BERT) (Reimers and Gurevych, 2019), and Scientific Text BERT (SciBERT) (Beltagy et al., 2019). These models were chosen due to their demonstrated capabilities in previous literature and practical usage in other Natural Language Processing (NLP) tasks, particularly in contexts requiring precise semantic understanding.

### Evaluation Dataset

The NFCorpus dataset^6^ from the BEIR framework was chosen for evaluation purposes. NFCorpus is specifically designed for information retrieval tasks, where the objective is to retrieve relevant documents based on natural language queries. It consists of over 3,000 fact-based documents, primarily drawn from the biomedical domain. Each query is paired with a set of relevance judgments, enabling the use of performance metrics such as F1-score, which requires ground truth labels for accurate evaluation. This makes NFCorpus an ideal benchmark for assessing how well a system understands and retrieves meaningful information, especially in the context of Quanta’s usage. Strong performance on this dataset suggests that the retrieval model is capable of handling complex, real-world queries - translating to more accurate and contextually relevant responses in chatbot applications (i.e. Quanta). Moreover, BEIR is known to provide a more standardized format (including documents, queries, and relevance labels), supporting reproducibility and fair comparison across retrieval methods.

### Evaluation Metrics

We employed five key performance metrics to comprehensively assess embedding model performance (**Table 3**).

**Table 3.**

| Metric | Description |
| --- | --- |
| **F1-score** | Measures the harmonic mean of precision and recall, indicating the balance between accuracy and completeness in retrieved results. |
| **Normalized Discounted Cumulative Gain (NDCG)** | Assesses the quality of retrieval results based on the position of relevant items, with higher-ranked correct items yielding a higher score. |
| **Recall@5** | Evaluates the proportion of relevant documents correctly retrieved within the top 5 results, emphasizing quick access to pertinent information. |
| **Mean Reciprocal Rank (MRR)** | Focuses on the rank position of the first relevant document retrieved, reflecting the retrieval system's ability to prioritize the most relevant information promptly. |
| **Computational Time** | Measures the speed at which each embedder processes queries and retrieves information, highlighting practical efficiency in real-world applications. |

Furthermore, to evaluate the stability, in addition to the score calculated, we’ve recorded the standard deviation of each embedder’s performance across all metrics mentioned above. A lower standard deviation indicates greater stability and consistent performance across various scenarios.

Prior to testing the effectiveness of LangChain, the requirements of the paired t-test were verified. Specifically, the observations were dependent (paired across the same query instances), and the normality of the paired differences was assessed using the Shapiro-Wilk test. While most metrics satisfied the normality assumption (p > 0.05), the MRR differences did not, with a Shapiro-Wilk p-value of 0.013. Therefore, F1 (p = 0.281), NDCG (p = 0.095), and Recall@5 (p = 0.069) were analyzed using the paired t-test, and non-parametric Wilcoxon signed-rank test was used for the MRR comparison to ensure statistical validity.

### Embedding Model Analysis

The evaluation procedure involved the following sequential steps. First, NFCorpus data was downloaded and limited to 500 documents and 20 queries for computational feasibility. This number, however, can be adjusted; the consequence of increasing the dataset is the increase in compile time. Then, each embedding model encoded the documents and queries into vector representations. After, both cosine and L2 similarities were computed using the FAISS library. For cosine similarity, all embeddings were normalized to ensure that the similarity score (ranges from -1 (completely dissimilar) to 1 (highly similar), with 0 indicating orthogonality) was based solely on vector direction rather than magnitude, as cosine similarity assumes unit-length vectors. The means and standard deviations of the 5 metrics were calculated and recorded in a CSV file for each embedding model with each similarity approach (cosine vs. L2). This CSV file was later used to graph a comparison bar graph of the means, as well as a heatmap for the standard deviation of each score.

### LangChain Evaluation & Integration

Upon identifying the optimal embedding model, LangChain was integrated into the architecture of Quanta to assess its potential for enhancement in semantic matching performance.

The process began by loading the NFCorpus dataset, limited to 500 documents and 50 queries to ensure efficiency during testing. The identified optimal embedding model (NLI-BERT) was then used to generate document embeddings, which were stored in a Facebook AI Similarity Search (FAISS) vector index to enable efficient similarity-based retrieval. A FakeListLLM was employed to simulate the language model component within the LangChain retrieval chain. This allowed us to isolate and evaluate the performance of the retrieval mechanism itself, without introducing variability from generated outputs and additional API cost. The LangChain retrieval chain combined the FAISS retriever with a custom “stuff” document chain to aggregate and format the top 5 most similar documents based on cosine similarity.

To assess LangChain’s impact, query-level retrieval performance was measured using four metrics: F1, NDCG, MRR, and Recall@5. Memory usage was tracked using Python’s tracemalloc module, and latency was measured using the time module. To determine the statistical significance of observed improvements, paired statistical tests were performed. Before applying the paired t-test, key conditions were verified: Query-level performance scores were paired across identical query instances. Specifically, the two samples correspond to the performance scores of the baseline (without LangChain) and LangChain systems, evaluated on the same set of queries via query_id, and the normality of the paired differences was assessed using the Shapiro-Wilk test, calculated with the shapiro function from the scipy.stats package.

While F1, NDCG, and Recall@5 differences met the normality assumption (p > 0.05), MRR did not (p = 0.013). Consequently, the paired t-test (ttest_rel from scipy.stats) was applied to the first three metric, while the Wilcoxon signed-rank test (non-parametric) was used to evaluate MRR. Each test computed the mean difference between LangChain and baseline scores and compared it to the distribution expected under the null hypothesis (no performance improvement), yielding p-values that confirmed the statistical significance of LangChain’s improvements, and not just due to random chance.

## Footnotes

1 https://python.langchain.com/

2 https://openai.com/index/gpt-4/

3 https://streamlit.io/

4 https://github.com/beir-cellar/beir

5 https://python.langchain.com/docs/tutorials/summarization/

6 https://www.cl.uni-heidelberg.de/statnlpgroup/nfcorpus/

## Notes

### Competing Interest Statement

The authors have declared no competing interest.

