## Supplementary Notes for "Accelerating Insight Discovery in Large Biomedical Text with Scalable Processing Framework"

### Supplementary Note 1: System Prompt

You are a highly capable AI research assistant specializing in ACADEMIC WRITING, SCHOLARLY ANALYSIS, and RESEARCH SUPPORT.
Your task is to generate a comprehensive, accurate, and well-substantiated response to the query below, taking ALL context and details provided into consideration.
Your output should reflect a deep understanding of the topic, integrate both provided context and relevant background knowledge, and adhere to the highest standards of academic rigor.
DO NOT GIVE AMBIGUOUS OR VAGUE RESPONSES. BE SPECIFIC USING DETAILS (ex. text, number, source of paper, date, etc.) WHEN AVAILABLE.
DO NOT INCLUDE ANY META-COMMENTARY (e.g., "the context shows," "the query asks," "based on the provided information," etc.).

YOUR RESPONSE SHOULD FOLLOW THESE 4 PROTOCOLS TOO:
1. Context Analysis:
- Carefully analyze the provided context to understand its relevance to the query.
- Identify key themes, arguments, or data points that directly relate to the query.

2. Query Understanding:
- Ensure that the query is fully understood. If the query is ambiguous, identify potential interpretations and choose the most logical one based on the context.
- Address the query based on its scope – whether it requires a direct answer, an in-depth analysis, or a synthesis of the information.

3. Answer Construction:
- Formulate a clear, logically structured response using precise and formal academic language.
- The answer should read as a naturally articulated, stand-alone academic explanation - DO NOT INCLUDE ANY META-COMMENTARY (e.g., "the context shows," "the query asks," "based on the provided information," etc.).
- When sufficient information is available, present a detailed and well-reasoned explanation, incorporating specific insights, data points, and arguments where relevant.
- Be SPECIFIC with the answers, referring to even the smallest details in the context if relevant.
- In cases where critical information is missing, enhance the answer using relevant, domain-specific knowledge.
- Distinguish clearly when supplemental information originates from general scholarly understanding rather than the provided materials.

- Use definitions, classifications, or examples to clarify technical terms or complex concepts when appropriate.
- Use appropriate styling such as bullet points or numbered lists for clarity, especially when presenting multiple points or steps in an argument.

4. Exceptions and Clarifications:
- If the query or context contains sensitive or ethical considerations, handle these with appropriate sensitivity and discretion.
- In the case of highly technical or domain-specific subjects, clarify complex terms and concepts to ensure understanding.

### Supplementary Note 2: Calculation of Memory Usage

**Average Memory Usage (Baseline BERT):**

From Table 1, baseline memory values (in bytes) are,

- 3754374, (3730572 × 9)
- Average = (3754374 + 9 × 3730572) / 10 = **3732952.2 bytes**

**Average Memory Usage (LangChain):**

From Table 2, memory values (in bytes) are,

- 525838, 128249, 105855, 107519, 106922, 112043, 104288, 103182, 101180, 105599
- Average = (Summation of bytes for all 10 trials) / 10 = 1,100,675 / 10 = **110067.5 bytes**

**Calculate Percentage Reduction:**

Reduction % = (1 − 110067.5/3732952.2) × 100 = (1 − 0.0299) × 100 ≈ **96.98%**

### Supplementary Note 3: Real-World Application

*Does prurigo nodularis have a genetic component?*

Prurigo nodularis (PN), a chronic inflammatory skin disease characterized by pruritic skin nodules, has an unknown etiology. The current understanding of the pathogenesis of PN is limited, with scant information available on genetic alterations that may contribute to the disease. The context does indicate that there may be a potential role for somatic events, which are often implicated in inflammatory conditions, in the development of PN.

Somatic events refer to genetic alterations that are acquired during an individual's lifetime, as opposed to germline mutations that are inherited. These acquired genetic alterations can lead to various diseases, including inflammatory conditions. In the case of PN, if somatic events play a role in its pathogenesis, this would suggest a potential genetic component to the disease. Furthermore, the observation that PN disproportionately affects African Americans could suggest a potential genetic or hereditary component, as this disparity may be reflective of genetic differences between populations. However, it is also possible that this disparity is due to non-genetic factors, such as environmental exposures or access to healthcare.

Given the current limited knowledge about the genetic underpinnings of PN, it cannot be conclusively stated that there is a genetic component to the disease. Further research, particularly studies investigating the genomic landscape and potential somatic alterations in individuals with PN, is necessary to elucidate the potential genetic contributions to the disease. It is also crucial to investigate the potential role of non-genetic factors in the observed disparities in PN prevalence among different populations.

*What comorbidities are associated with higher mortality in PN patients, and how do these vary by race?*

PN is a chronic inflammatory skin condition characterized by intensely pruritic and hyperkeratotic nodules, predominantly on the torso and extremities. The available evidence shows that PN is associated with a high burden of systemic and psychiatric comorbidities, and disproportionately affects racial and ethnic minorities.

A large-scale multi-center cohort study using TriNetX data (Sutaria et al., 2022) found that PN patients experience significantly higher all-cause mortality than matched controls (Hazard Ratio [HR] 1.70). When stratified by race, Black patients had the highest mortality risk (HR 2.07), followed by White (HR 1.74) and Hispanic patients (HR 1.62). Interestingly, increased mortality was not observed among Asian PN patients. These disparities persist even after adjusting for age, sex, and other baseline demographic factors.

While the specific comorbidities directly contributing to increased mortality in PN were not exhaustively listed in that study, PN has been previously linked to several conditions, including:

- Chronic kidney disease
- Diabetes
- Malignancies
- Cardiovascular disease
- Depression

The elevated mortality in PN patients likely reflects the combined impact of these chronic illnesses. Moreover, recent studies suggest that Black PN patients may exhibit stronger systemic inflammatory responses, particularly involving Th22 cytokine pathways, which could contribute to worse outcomes.

The reasons behind these racial disparities are likely multifactorial, involving differences in comorbid disease prevalence, healthcare access, environmental exposures, and possibly genetic susceptibility.
In summary, while PN is clearly associated with increased mortality - especially among Black patients - the underlying contributors appear to include a combination of comorbid conditions and systemic inflammation, with racial disparities pointing to broader socio-biological factors that warrant further investigation.

*Which neural receptors (e.g., TRPV1, Substance P pathways) are differentially expressed in pruritic vs. non-pruritic skin?*
Two neural receptors, TRPV1 (Transient Receptor Potential Vanilloid 1) and TRPA1 (Transient Receptor Potential Ankyrin 1), exhibit variations in expression in pruritic (itchy) and non-pruritic skin, as highlighted in the context.
TRPV1 (Transient Receptor Potential Vanilloid 1) is significantly overexpressed in pruritic lesional skin compared to non-pruritic skin in both AD and psoriasis. TRPV1, which is activated by stimuli such as heat and capsaicin, plays a key role in pain and itch signaling. Its expression levels were found to correlate positively with itch intensity, implicating it as a major neural mediator of pruritus.
TRPA1 (TRP Ankyrin 1), another TRP family member, is overexpressed specifically in pruritic atopic skin, but not in non-pruritic areas. TRPA1 is known to respond to environmental irritants and oxidative stress, and contributes to itch through activation of sensory neurons.
Substance P (encoded by TAC1) and its receptor Neurokinin-1 (TACR1) were also significantly upregulated in pruritic skin, supporting their role in mediating neurogenic inflammation and pruritus. Substance P is a neuropeptide that can induce mast cell degranulation, vasodilation, and direct activation of sensory nerves.
In contrast, non-pruritic skin from the same patients typically showed lower or baseline expression levels of these receptors.
These findings indicate that specific neural receptors are upregulated in itchy skin lesions, and many of them correlate with subjective itch scores, suggesting a mechanistic role in chronic pruritus. While overexpression does not prove causality, these targets offer promising avenues for therapeutic intervention in itch-dominant dermatologic diseases.

*What are the limitations or potential drawbacks of the research described in the papers?*
The limitations or potential drawbacks of the research mentioned predominantly revolve around data quality and availability, as well as the time and financial resources required to conduct the study.

1. Data Quality: The research is heavily dependent on the availability of high-quality data. If the data is inaccurate, incomplete, or outdated, it could lead to errors in the final output. Similarly, if the data is not representative of the population or area being studied, it could lead to bias.
2. Time and Cost: The creation of realistic maps is an elaborate process that takes considerably longer to complete than conventional maps. This could make the research time-consuming and costly, which might be a significant drawback if resources are limited or if rapid results are needed.
3. Funder Influence: The research mentions that the production of realistic maps, a significant component of the study, is an elaborate process that takes considerably longer to complete than their conventional counterparts. This implies that the research process could be time-consuming, which might delay the attainment of results and potentially increase the cost of the project. In addition, it might also limit the ability of researchers to respond quickly to changes or advancements in the field during the course of the study.
